# Circulating Sex-specific Markers of Plaque Instability in Women and Men with Severe Carotid Atherosclerosis

**DOI:** 10.1101/2023.06.12.544697

**Authors:** Karina Gasbarrino, Huaien Zheng, Stella S. Daskalopoulou

## Abstract

**Background:** Differences in plaque composition and instability exist among men and women. Circulating markers that reflect sex-specific features in the plaque should be explored for better prediction of high-risk plaques in women and in men. This study aims to 1) investigate differences in the lipid, immune, and adipokine circulating profiles of men and women with stable versus unstable plaques, and 2) identify circulating markers that can better classify men and women according to plaque instability.

**Methods:** Pre-operative blood samples as well as plaque specimens were collected from men and women undergoing a carotid endarterectomy (n=460). Blood samples were used for adipokine, lipid, and immune profiling. Plaque stability was determined by gold-standard histological classifications.

**Results:** Men had more unstable plaques than women (P<0.001), exhibiting greater plaque hemorrhage, a larger lipid core, and more inflammation (P<0.001), as well as less favourable circulating profiles. Significant antagonistic interactions were observed between sex and white blood cell (WBC) counts, sex and basophil to WBC ratio, and sex and platelet counts on impacting plaque instability. Several circulating immune parameters served as independent sex-specific markers of plaque instability; low total white blood cell (WBC) counts, high monocyte to WBC ratio, and low basophil to WBC ratio were associated with greater plaque instability in men, while a higher basophil to WBC ratio was observed in women with unstable plaques.

**Conclusions:** Our findings demonstrated sex-specific differences between older men and postmenopausal women with severe carotid atherosclerosis, with women displaying more stable plaque phenotypes, and favourable circulating profiles compared to men. We identified several potential circulating markers that relate to sex-specific plaque phenotypes for better prediction of high-risk plaques in women and in men. Following future validation, these markers could be implemented into clinical practice to monitor when the plaque becomes unstable and better select men and women for intervention.

## 1. Introduction

Stroke remains one of the leading causes of mortality and disability worldwide^1^. Although stroke incidence rates are higher among men than women, it has been reported that women have worse recovery and higher mortality and disability rates than men post-stroke^2–4^. Most ischemic strokes are caused by cerebral ischemia, with a major contributing factor being the rupture of unstable atherosclerotic plaques that form in the internal carotid artery. Unstable plaques are characterized by a large lipid-rich core, a thin fibrous cap, ulceration, thrombosis, and intraplaque hemorrhage, as well as an abundance of inflammatory cells^5^. It is well established that differences in plaque morphology and composition exist among men and women, where men are more likely to develop unstable and complex plaques with more inflammatory features than women, while women’s plaques are characterized by more stable histological features and higher-grade stenoses^6–8^.

Although the above-mentioned sex disparities have led to an increased interest in understanding the mechanisms underlying the atherosclerotic process in women versus men, gaps in optimal clinical practice still exist, such as the lack of sex-specific guidelines for carotid atherosclerotic disease management. Current guidelines recommend a carotid endarterectomy (CEA) for stroke prevention in men and women with symptomatic and asymptomatic moderate-to-high-grade carotid artery stenosis^9, 10^. However, these guidelines were based off carotid stenosis trials where women were under-represented, and the evidence was mainly extrapolated from research performed in men. In fact, further research suggests that women benefit less from carotid revascularization than men and have an increased surgical risk^7, 10–12^. Differences in plaque morphology and composition are believed to partly explain why women benefit less from surgical intervention than men^7^. Thus, there is a need to identify circulating markers that reflect sex-specific features in the plaque for better prediction of high-risk plaques in women and in men. The aim of this study is two-fold: 1) to investigate differences in the lipid, immune, and adipokine circulating profiles of men and women with stable versus unstable atherosclerotic plaques, and 2) identify circulating markers that can better classify men and women according to plaque instability. A better understanding of these differences between men and women with severe carotid atherosclerosis and the identification of potential circulating markers will allow for more appropriate and sex-specific prevention, diagnosis, and treatment measures.

## 2. Methods

### 2.1 Study population

Consecutive eligible, neurologically symptomatic (defined as having symptoms within 6 months prior to surgical intervention), and asymptomatic men and women, undergoing a CEA due to clinical indications, were recruited from preoperative clinics at the McGill University Health Centre (MUHC) and the Jewish General Hospital (JGH), as described previously^13, 14^. In the preoperative clinics, extensive assessments were performed to rule out non-carotid causes of cerebrovascular events in symptomatic patients. Our study complies with the Declaration of Helsinki. Ethics approval has been granted by the McGill University’s Institutional Ethics Review Board (MP-37-2018-3789), and all study participants provided written informed consent. Excluded from this study were premenopausal women, men or women who are taking/or have taken hormone replacement therapy, as well as patients with previous interventions on the same carotid artery (CEA or carotid stent), as the histopathology of the re-stenotic plaque may be altered^15, 16^. Postmenopausal women were defined as women who reported absence of menstrual periods for at least 12 months.

Sociodemographic and clinical patient information (i.e., cerebrovascular symptomatic status, carotid artery stenosis, past medical history, medication use, and lifestyle habits) were obtained and cross-matched through various sources: 1) patient interview, 2) a detailed questionnaire, and 3) medical records. Blood pressure measurements were obtained according to guidelines^17^, and anthropometric measurements (height, weight, body mass index [BMI]) were collected using standardized methods.

### 2.2 Blood collection and measurements

Blood sample collection and analysis methods have been previously reported^13, 14^. Fasting blood samples were collected from each subject preoperatively on the morning of the CEA and were used to obtain plasma and serum for subsequent biochemical and hormone analyses.

#### Circulating Lipid, Immune, and Adipokine Analyses

Serum lipid profile (triglycerides, total cholesterol, high-density lipoprotein cholesterol [HDL-C], apolipoprotein A1 [apoA1], and apolipoprotein B [apoB]), high-sensitivity C-reactive protein (hsCRP), fasting glucose levels, and complete blood count (including white blood cell count [WBC], red blood cell count, hemoglobin and hematocrit levels, and platelet count) were measured at the MUHC central biochemistry lab, using standardized methods. Low-density lipoprotein cholesterol (LDL-C) levels were calculated using the Friedewald formula^18^. Non-HDL-C and remnant cholesterol levels were calculated using the following formulas, respectively: 1) total cholesterol – HDL-C, 2) total cholesterol - (LDL-C + HDL-C).

Profiling of collected sera for the simultaneous quantification of selected cytokine, chemokine, and vascular injury markers (interferon [IFN]-γ, interleukin [IL]-1β, IL-6, IL-10, macrophage inflammatory protein [MIP]-1α, tumor necrosis factor [TNF]-α) was performed in a representative subset of our participants using an adapted version of the MILLIPLEX^®^ Human High Sensitivity T Cell Magnetic Bead Panel (Millipore Sigma, Burlington, MA, USA). The MAGPIX^®^ Luminex analyzer (Luminex Corporation, Austin, TX, USA) was used to acquire and analyze the data. Intra-assay coefficients of variation were <5% and inter-assay coefficients of variation were <20%.

Plasma chemerin, resistin, and total and high molecular weight (HMW) adiponectin levels were measured by enzyme-linked immunosorbent assays (Human Chemerin ELISA kit, Biovendor, Brno, Czech Republic; Human Resistin ELISA kit, Millipore Sigma; Human Total Adiponectin/Acrp30 Quantikine ELISA Kit, R&D Systems, Minneapolis, MN, USA; Human HMW Adiponectin/Acrp30 Quantikine ELISA Kit, R&D Systems). All samples were run in duplicate, and the lower limits of detection were 0.1 ng/mL for chemerin, 0.02 ng/mL for resistin, and 0.891 ng/mL and 0.989 ng/mL for total and HMW adiponectin, respectively. Inter-and intra-assay coefficients of variation were <10%. Please see the Major Resources Table in the Supplemental Materials.

### 2.3 Histological classification of carotid atherosclerotic plaques

Carotid plaque specimens were obtained immediately following surgical resection and were processed for histological analyses, as previously described^13, 14, 19, 20^. Four micrometer sections from the plaque segment with the area of maximal stenosis and largest plaque burden were stained with hematoxylin and eosin, and with antibodies against markers to identify lymphocytes, macrophages, smooth muscle cells, and neovessels (CD3, CD68, α-SMC actin, von Willebrand factor, respectively). Plaque composition and stability were characterized by vascular pathologists (Drs. Veinot and Lai), according to two gold-standard classifications of plaque instability: 1) American Heart Association (AHA) classification by Stary *et al*.^21^, and 2) semi-quantitative scale by Lovett *et al*^22, 23^, as previously described^13, 14^. The pathologists were unaware of patients’ cerebrovascular symptomatic status and all demographic and clinical information.

### 2.4 Statistical analyses

Chi-square (χ2), analysis of variance (parametric test) or Kruskal-Wallis test (non-parametric test), were used as appropriate, to assess the differences in clinical characteristics, cerebrovascular symptomatology, histological features of the plaque, circulating lipid, immune, and adipokine markers, and blood count parameters between men and women with stable and unstable carotid plaques. The definitely stable and probably stable plaque groups, classified by the vascular pathologists according to Lovett’s semi-quantitative scale, were combined and labeled as ‘stable’, while the probably unstable and definitely unstable plaque groups were combined and labeled as ‘unstable’ to be used in the statistical analyses. In addition to being clinically relevant, combining instability groups led to increased statistical power.

Logistic regression analyses were performed to estimate : 1) the effect of sex on plaque instability (adjustments for age, BMI, history of coronary artery disease [CAD], type 2 diabetes mellitus [T2DM], hypertension, carotid artery stenosis, cerebrovascular symptomatic status, statin use, WBCs, platelet counts, HMW to total adiponectin ratio, hsCRP, and LDL-C levels), 2) the effect of the interaction between sex and various lipid, immune, and adipokine parameters on plaque instability (Model 1: adjusted for age and BMI; Model 2: Model 1 + adjusted for history of CAD, T2DM, hypertension, carotid artery stenosis, cerebrovascular symptomatic status, statin use, hsCRP; Model 3: Model 2 + LDL-C, WBCs, platelet counts, and HMW to total adiponectin ratio, where appropriate), and 3) the sex-specific effect of various lipid, immune, and adipokine parameters on plaque instability (same models tested as in 2). Odd ratios (ORs) are presented with 95% confidence intervals (CIs). All statistical analyses were performed in SPSS, Version 20 (IBM, Armonk, New York, United States). Values of P<0.05 (2-tailed) were considered significant.

## 3. Results

### 3.1 Sex differences in clinical characteristics in relation to plaque instability

In our study population over twice as many men underwent a CEA than women (31.1% women/68.9% men). **Table 1** summarizes the baseline clinical characteristics of men and women according to plaque stability. Age, BMI, and blood pressure were similar among the four patient groups (women with stable plaques, men with stable plaques, women with unstable plaques, men with unstable plaques). Both men and women suffered from a high presence of comorbidities, such as hypertension, hypercholesterolemia, and T2DM. Medication use (statins, anti-hypertensive, anti-hyperglycemic, and anti-platelet medication) did not differ between men and women.

**Table 1.** Sex-specific differences in patient clinical characteristics in relation to plaque instability.

| Clinical Characteristics | Women (n=143) |  | Men (n=317) |  | P-value |
| --- | --- | --- | --- | --- | --- |
|  | With Stable Plaques (n=84) | With Unstable Plaques (n=59) | With Stable Plaques (n=122) | With Unstable Plaques (n=195) |  |
| Age, y | 71.8±8.1 | 70.0±9.5 | 71.0±9.1 | 70.9±9.3 | 0.690 |
| BMI, kg/m<sup>2</sup> | 27.8±5.2 | 26.3±4.4 | 26.7±3.7 | 27.5±4.5 | 0.200 |
| Ever smoker, % | 65.1 | 73.7 | 71.9 | 78.8 | 0.114 |
| CAD, % | 32.1 | 28.8 | 46.7 | 41.0 | 0.056 |
| Cerebrovascular Symptomatic Status, % | 79.8 | 84.7 | 72.1 | 79.5 | 0.218 |
| Carotid Artery Stenosis, 50-79%/80-99% | 29.6/70.4 | 22.0/78.0 | 37.0/63.0 | 30.5/69.5 | 0.231 |
| SBP, mmHg | 141±22 | 137±21 | 138±18 | 138±19 | 0.743 |
| DBP, mmHg | 72±10 | 72±12 | 73±10 | 74±10 | 0.483 |
| Hypertension, % | 82.1 | 84.7 | 88.5 | 83.1 | 0.537 |
| Anti-hypertensive medication, % | 95.7 | 94.0 | 96.3 | 92.6 | 0.580 |
| Glucose, mmol/L | 6.00 [5.30-7.26] | 6.36 [5.50-7.66] | 5.90 [5.30-7.25] | 6.38 [5.50-7.66] | 0.199 |
| T2DM, % | 32.5 | 23.7 | 39.3 | 33.3 | 0.218 |
| Anti-hyperglycemic medication, % | 100.0 | 92.9 | 95.8 | 87.7 | 0.149 |
| Hypercholesterolemia, % | 79.8 | 84.7 | 84.4 | 82.1 | 0.805 |
| Statin use, % | 76.2 | 83.1 | 86.1 | 74.4 | 0.067 |
| Anti-platelets, % | 92.8 | 90.9 | 96.1 | 89.3 | 0.255 |

Values are either represented as mean ± standard deviation or median [interquartile range] (for continuous data) or as percentages (for nominal data).
BMI indicates body mass index; CAD, coronary artery disease; SBP, systolic blood pressure; DBP, diastolic blood pressure; T2DM, type 2 diabetes mellitus.
P-value indicates comparison among all groups: women with stable plaques, women with unstable plaques, men with stable plaques, men with unstable plaques (analysis was performed by one-way ANOVA, Kruskal-Wallis, or Chi-square ( $\chi^2$ ) test, as appropriate).

### 3.2 Sex differences in plaque instability and clinical presentation

According to Lovett’s semi-quantitative scale of plaque instability, a greater proportion of women had definitely stable and probably stable plaques, while a greater proportion of men had definitely unstable and probably unstable plaques (P<0.001; **Figure 1A**). Classifying plaques according to the AHA histological classification further confirmed these results; women had a significantly greater proportion of Type VII and VIII plaques than men, which were composed predominantly of calcification and fibrous tissue, respectively. On the other hand, men had a greater proportion of Type V and Type VI plaques than women, which are characterized as more complicated and unstable lesions (P<0.001; **Figure 1B**). Univariate logistic regression analyses demonstrated that being a man increased the odds of having an unstable plaque by 128% (OR [95% CI]: 2.28 [1.52-3.40]; P<0.001) compared to women. This association remained significant after adjustment for age, BMI, history of CAD, T2DM, hypertension, carotid artery stenosis, cerebrovascular symptomatic status, statin use, WBCs, platelet counts, HMW to total adiponectin ratio, cortisol, hsCRP, and LDL-C levels (2.06 [1.24-3.43]; P=0.006). Looking at the individual features of the plaque, men’s plaques were significantly characterized by more unstable histological features than women, such as a larger hemorrhage, less fibrous tissue, more foam cells, a larger lipid core, and more inflammation (macrophage and lymphocyte infiltration) overall in the plaque as well as in the fibrous cap of the plaque (**Table 2**). However, despite these differences in plaque composition, a similar proportion of men and women were symptomatic and experienced similar types of ischemic events (**Figure 1C-D**). Interestingly, among those individuals who were symptomatic, significant differences in plaque composition were still observed between men and women; men’s plaques were characterized by a larger hemorrhage (P=0.034) and lipid core (P<0.001), less fibrous tissue (P<0.001), and a greater presence of foam cells (P=0.009) and inflammation (P<0.01), and cap infiltration (P=0.002) than women. In contrast, the presence of a thrombus or ruptured cap was similar between men and women who experienced a symptomatic event (P>0.05).

**Figure 1.**
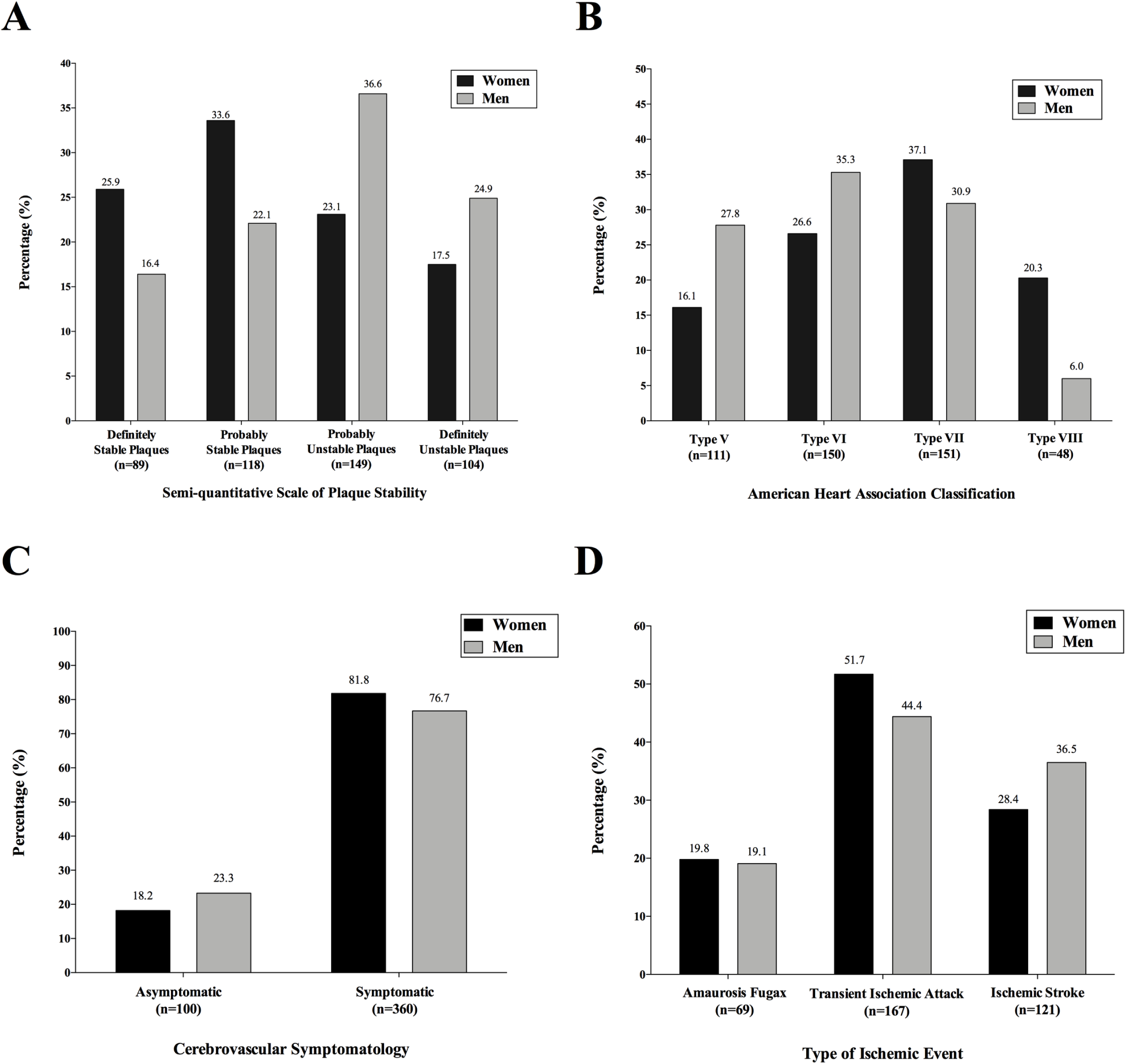
Sex-specific differences in plaque stability and cerebrovascular symptomatology in CEA population. Bar graphs represent the percentage of **A)** women and men with stable or unstable plaques according to Lovett’s semi-quantitative scale of plaque stability, P<0.001 (Chi-square test), **B)** women and men with Type V, VI, VII, or VIII plaques according to the AHA classification, P<0.001 (Chi-square test), **C)** women and men who were symptomatic or asymptomatic, P=0.214 (Chi-square test), **D)** symptomatic women and men according to the type of cerebrovascular event they suffered, P=0.298 (Chi-square test).

**Table 2.**
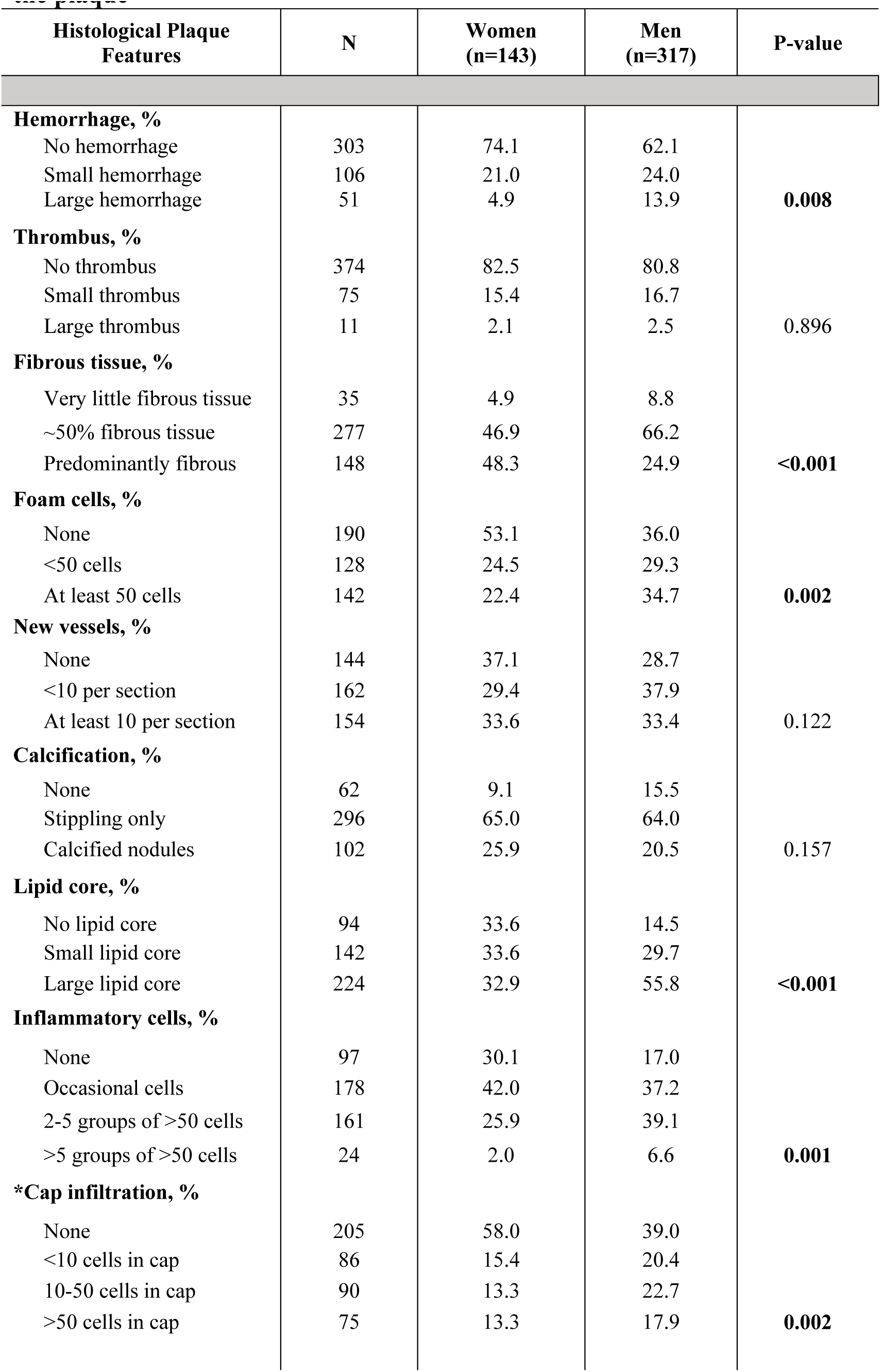
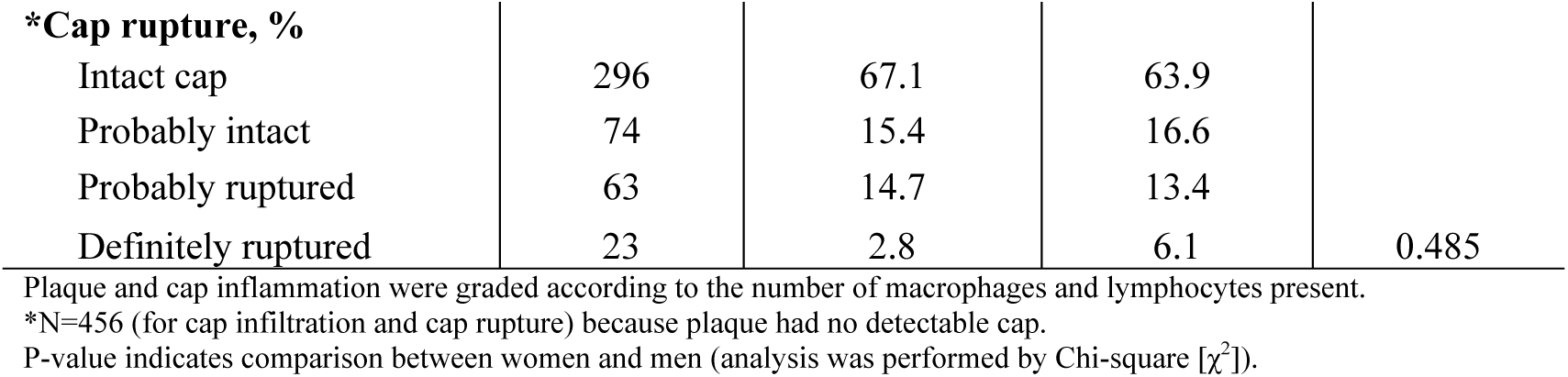
Sex-specific differences in the prevalence of histological features of the plaque.

### 3.3 Sex differences in circulating lipid, immune, and adipokine markers in relation to plaque instability

**Table 3** summarizes the differences in lipid, immune, and adipokine markers, and blood count parameters between men and women with stable versus unstable plaques. HDL-C levels were significantly higher in women compared with men, irrespective of plaque stability (**Table 3**). As a result, apoA1 levels were also significantly higher among women compared to men whether plaques were classified as stable or unstable (**Table 3**). The total circulating WBC count was observed to be significantly higher in men versus women when plaques were stable (P=0.003; **Table 3**). Interestingly, the total WBC count was significantly decreased in men with unstable vs stable plaques (P=0.001), along with the proportion of basophil to WBC count (P=0.001; **Table 3**). Men with unstable plaques had the lowest proportion of circulating lymphocyte to WBC count (P=0.002), and the greatest proportion of circulating monocytes to WBC count (P=0.006; **Table 3**), while the opposite was observed in women with stable plaques. However, no difference in circulating levels of cytokines and chemokines, hsCRP, IL-6, TNF-α, MIP1-α, IL-1β, IFN-γ, and IL-10 were detected among the patient groups (**Table 3**). The total red blood cell count, along with hemoglobin and hematocrit levels were significantly higher in men compared with women, irrespective of plaque stability (**Table 3**). The platelet count, on the other hand, was significantly higher among women with unstable plaques compared with men with unstable plaques (P<0.001) but remained similar among men and women with stable plaques (**Table 3**). Men with unstable plaques had significantly lower total platelet counts versus men with stable plaques (P=0.016), while no significant differences were observed among women with unstable versus stable plaques (**Table 3**). Women had significantly higher total and HMW adiponectin levels compared with men, irrespective of plaque stability (**Table 3**). However, the proportion of HMW to total adiponectin levels was only significantly different between women and men with stable plaques (P<0.001), and not between women and men who had unstable plaques (**Table 3**).

**Table 3.**
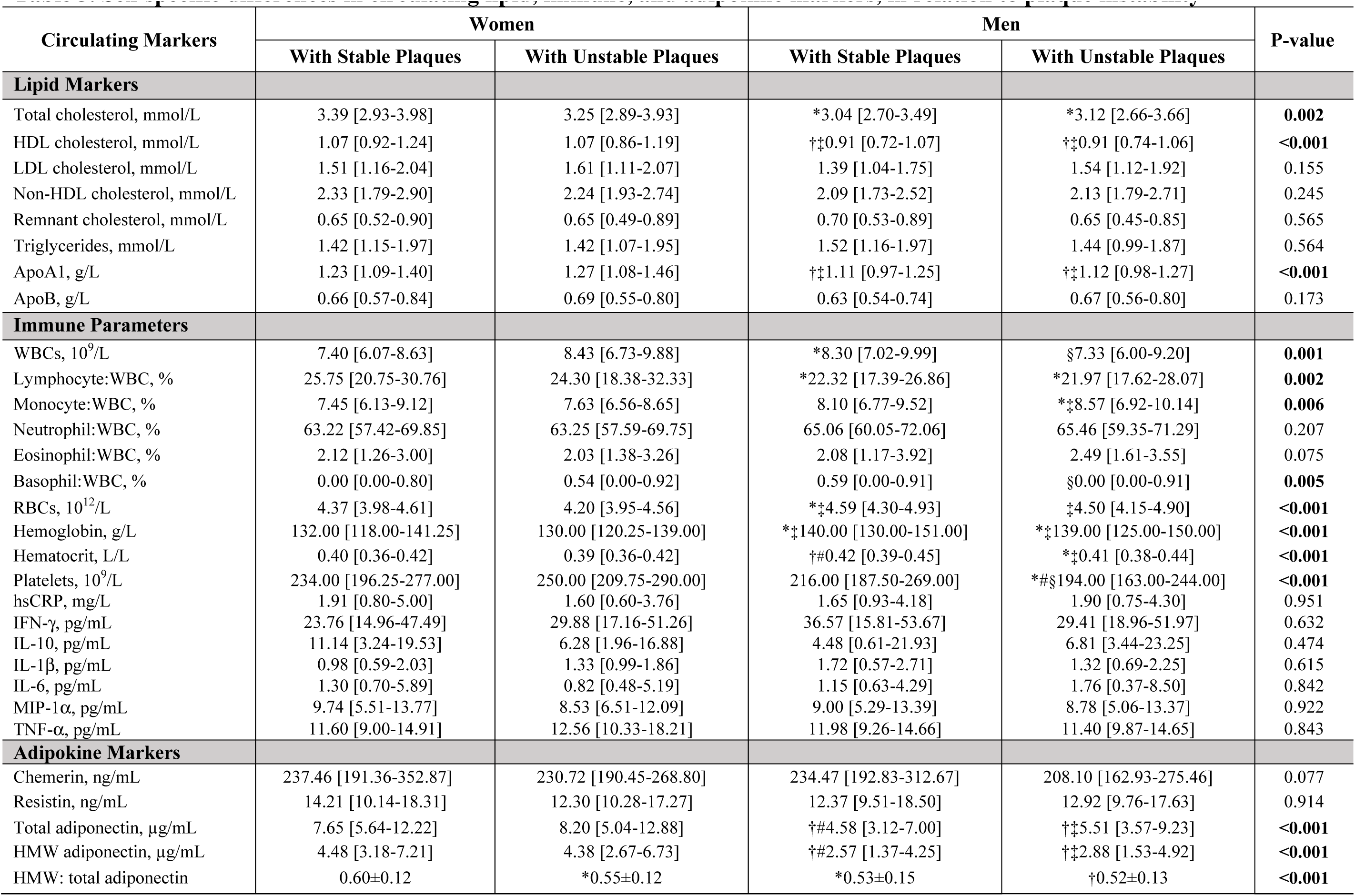
Sex-specific differences in circulating lipid, immune, and adipokine markers, in relation to plaque instability.

This may be due to the fact that women with unstable plaques had significantly lower HMW:total adiponectin than women with stable plaques (P=0.034, **Table 3**).

Interaction regression models were performed to determine the presence of interaction between biological sex and various lipid, immune, and adipokine markers, and plaque instability. We observed significant interactions between sex and WBC counts, sex and basophil to WBC ratio, and sex and platelet counts on impacting plaque instability (**Table 4**). In the fully adjusted model (Model 3), male sex, a higher WBC count, or a higher basophil to WBC ratio was significantly and independently associated with greater plaque instability, while platelet counts alone had no effect on plaque instability (**Table 4**). However, an antagonistic interaction was observed in males in the presence of high WBC counts, a high basophil to WBC ratio, or high platelet counts (**Table 4**).

**Table 4.** Interaction Logistic Regression Models using Plaque Instability as a Dependent Variable.

|  | Univariate |  | Model 1 |  | Model 2 |  | Model 3 |  |
| --- | --- | --- | --- | --- | --- | --- | --- | --- |
| | $\beta$<br>(95% CI) | P-value | $\beta$<br>(95% CI) | P-value | $\beta$<br>(95% CI) | P-value | $\beta$<br>(95% CI) | P-value |
| **Sex\*Low-density lipoprotein cholesterol** | 1.307 (0.868-1.967) | 0.199 | 1.297 (0.842-1.998) | 0.238 | 1.470 (0.932-2.317) | 0.097 | 1.434 (0.877-2.344) | 0.151 |
| Sex | **2.426 (1.609-3.657)** | **<0.001** | **1.920 (1.241-2.968)** | **0.003** | **1.990 (1.258-3.147)** | **0.003** | **1.979 (1.205-3.249)** | **0.007** |
| **Low-density lipoprotein cholesterol** | 1.025 (0.754-1.392) | 0.876 | 1.059 (0.767-1.462) | 0.728 | 0.979 (0.694-1.381) | 0.905 | 1.001 (0.685-1.464) | 0.995 |
| **Sex\*White blood cell counts** | **0.619 (0.408-0.940)** | **0.024** | 0.679 (0.433-1.064) | 0.091 | 0.621 (0.385-1.001) | 0.051 | **0.275 (0.136-0.558)** | **<0.001** |
| Sex | **2.294 (1.529-3.441)** | **<0.001** | **1.823 (1.185-2.805)** | **0.006** | **1.936 (1.225-3.060)** | **0.005** | **2.849 (1.649-4.923)** | **<0.001** |
| **White blood cell counts** | 1.270 (0.900-1.792) | 0.173 | 1.155 (0.800-1.667) | 0.442 | 1.202 (0.812-1.780) | 0.358 | **2.026 (1.129-3.636)** | **0.018** |
| **Sex\*Monocyte: white blood cell** | 1.331 (0.821-2.160) | 0.247 | 1.154 (0.674-1.975) | 0.601 | 1.161 (0.665-2.026) | 0.600 | 1.185 (0.667-2.106) | 0.562 |
| Sex | **2.285 (1.483-3.521)** | **<0.001** | **1.804 (1.137-2.863)** | **0.012** | **1.925 (1.181-3.138)** | **0.009** | **1.837 (1.103-3.058)** | **0.019** |
| **Monocyte: white blood cell** | 1.019 (0.680-1.528) | 0.927 | 1.229 (0.781-1.932) | 0.373 | 1.196 (0.746-1.917) | 0.457 | 1.236 (0.757-2.018) | 0.396 |
| **Sex\*Basophil: white blood cell** | **0.627 (0.415-0.949)** | **0.027** | **0.467 (0.283-0.772)** | **0.003** | **0.369 (0.215-0.635)** | **<0.001** | **0.373 (0.215-0.647)** | **<0.001** |
| Sex | **2.347 (1.532-3.595)** | **<0.001** | **1.902 (1.206-3.000)** | **0.006** | **2.041 (1.257-3.314)** | **0.004** | **1.969 (1.188-3.263)** | **0.009** |
| **Basophil: white blood cell** | 1.174 (0.842-1.637) | 0.345 | 1.419 (0.938-2.146) | 0.097 | **1.759 (1.127-2.747)** | **0.013** | **1.806 (1.147-2.845)** | **0.011** |
| **Sex\*Platelet counts** | **0.595 (0.377-0.940)** | **0.026** | **0.565 (0.344-0.929)** | **0.024** | **0.541 (0.318-0.921)** | **0.024** | **0.532 (0.305-0.929)** | **0.026** |
| Sex | **2.411 (1.553-3.741)** | **<0.001** | **2.049 (1.282-3.277)** | **0.003** | **2.200 (1.341-3.608)** | **0.002** | **2.214 (1.328-3.694)** | **0.002** |
| **Platelet counts** | 1.273 (0.866-1.872) | 0.219 | 1.375 (0.898-2.106) | 0.142 | 1.407 (0.886-2.236) | 0.148 | 1.511 (0.912-2.505) | 0.109 |
| **Sex\*High-molecular weight: total adiponectin** | 1.453 (0.911-2.319) | 0.117 | 1.486 (0.887-2.491) | 0.133 | 1.362 (0.792-2.340) | 0.264 | 1.215 (0.686-2.152) | 0.504 |
| Sex | **2.028 (1.331-3.092)** | **0.001** | **1.634 (1.045-2.555)** | **0.031** | **1.681 (1.051-2.688)** | **0.030** | **1.914 (1.156-3.168)** | **0.012** |
| **High-molecular weight: total adiponectin** | **0.641 (0.425-0.967)** | **0.034** | 0.650 (0.413-1.021) | 0.062 | 0.632 (0.393-1.017) | 0.059 | 0.652 (0.398-1.068) | 0.090 |

Results were the associations of each unit increase in the levels of the variable of interest with plaque instability. Values are represented as odds ratio (95% confidence interval).
Model 1: adjusted for age and body mass index
Model 2: Model 1 + adjusted for history of coronary artery disease, type 2 diabetes mellitus, hypertension, carotid artery stenosis, cerebrovascular symptomatic status, statin use, high-sensitivity C-reactive protein
Model 3: Model 2 + low-density lipoprotein cholesterol, white blood cell counts, platelet counts, and high-molecular weight to total adiponectin ratio, where appropriate.
\*P<0.05; †P≤0.01; ‡P≤0.001

Multivariate logistic regression models were also performed to identify independent sex-specific markers of plaque instability (**Table 5**). In men, low total WBC counts, a high monocyte to WBC ratio, and a low basophil to WBC ratio were observed to be independently associated with increased plaque instability, even in the fully adjusted model (adjustments for age, BMI, history of CAD, T2DM, hypertension, carotid artery stenosis, cerebrovascular symptomatic status, statin use, cortisol, hsCRP, LDL-C, platelet counts, and HMW to total adiponectin ratio), while high LDL-C levels were directly associated with plaque instability only in Models 1 and 2 (**Table 5**). On the other hand, in the fully adjusted model, exposure to a high basophil to WBC ratio increased the odds of women having an unstable plaque by 214% (**Table 5**). Furthermore, a high HMW to total adiponectin ratio was significantly associated with decreased plaque instability in women only in Models 1 and 2 (**Table 5**).

**Table 5.** Logistic regression models of the association between lipid, immune, and adipokine markers and carotid plaque instability in men and women.

|  | **Men**<br>**OR (95% CI)** | **Women**<br>**OR (95% CI)** |
| --- | --- | --- |
| *Low-density lipoprotein cholesterol* |  |  |
| Univariate model | **\*1.485 (1.030-2.141)** | 1.034 (0.682-1.565) |
| Model 1 | **\*1.579 (1.069-2.332)** | 1.076 (0.686-1.688) |
| Model 2 | **\*1.580 (1.008-2.475)** | 1.127 (0.651-1.951) |
| Model 3 | 1.484 (0.929-2.372) | 0.902 (0.475-1.713) |
| *White blood cell counts* |  |  |
| Univariate model | **\*0.924 (0.854-0.999)** | 1.082 (0.966-1.213) |
| Model 1 | 0.925 (0.848-1.008) | 1.081 (0.952-1.228) |
| Model 2 | **\*0.911 (0.831-0.998)** | 1.078 (0.933-1.246) |
| Model 3 | **\*0.827 (0.713-0.960)** | 1.254 (0.992-1.584) |
| *Monocyte: white blood cell* |  |  |
| Univariate model | **\*1.116 (1.014-1.228)** | 1.007 (0.870-1.165) |
| Model 1 | **\*1.129 (1.017-1.253)** | 1.080 (0.914-1.276) |
| Model 2 | **\*1.135 (1.012-1.273)** | 1.055 (0.868-1.281) |
| Model 3 | **\*1.158 (1.027-1.305)** | 1.058 (0.868-1.288) |
| *Basophil: white blood cell* |  |  |
| Univariate model | **\*0.569 (0.362-0.895)** | 1.343 (0.728-2.476) |
| Model 1 | †**0.468 (0.279-0.785)** | 2.024 (0.928-4.416) |
| Model 2 | †**0.445 (0.256-0.773)** | **\*3.041 (1.211-7.634)** |
| Model 3 | **\*0.495 (0.281-0.871)** | **\*3.142 (1.220-8.093)** |
| *Platelet counts* |  |  |
| Univariate model | **\*0.996 (0.993-1.000)** | 1.003 (0.998-1.008) |
| Model 1 | 0.997 (0.993-1.000) | 1.005 (0.999-1.011) |
| Model 2 | 0.997 (0.993-1.000) | 1.005 (0.998-1.011) |
| Model 3 | 0.998 (0.994-1.002) | 1.002 (0.994-1.010) |
| *High-molecular weight: total adiponectin* |  |  |
| Univariate model | 0.595 (0.118-3.011) | **\*0.039 (0.002-0.782)** |
| Model 1 | 0.853 (0.135-5.403) | **\*0.031 (0.001-0.939)** |
| Model 2 | 0.398 (0.053-2.989) | **\*0.013 (0.000-0.613)** |
| Model 3 | 0.209 (0.022-1.960) | 0.024 (0.000-1.395) |

Results were the associations of each unit increase in the levels of the variable of interest with plaque instability. Values are represented as odds ratio (95% confidence interval).
Model 1: adjusted for age and body mass index
Model 2: Model 1 + adjusted for history of coronary artery disease, type 2 diabetes mellitus, hypertension, carotid artery stenosis, cerebrovascular symptomatic status, statin use, high-sensitivity C-reactive protein
Model 3: Model 2 + low-density lipoprotein cholesterol, white blood cell counts, platelet counts, and high-molecular weight to total adiponectin ratio, where appropriate.
\*P<0.05; †P≤0.01; ‡P≤0.001

## 4. Discussion

Our study provided a comprehensive characterization of sex differences that exist between older men and postmenopausal women with severe carotid atherosclerosis who underwent a CEA based on the current guidelines for carotid disease management. We demonstrated that men and women exhibited clear differences not only at the level of the plaque but also at the level of the circulation, with women displaying more favourable adipokine, lipid, and immune profiles compared to men, often irrespective of the stability of their plaque and despite being post-menopausal for many years (mean age: 71 years). Interestingly, an antagonistic interaction was revealed between biological male sex and various circulating parameters (WBC counts, basophil to WBC ratio, platelet counts) on influencing plaque instability. Several immune parameters were also identified to serve as independent sex-specific markers of plaque instability.

### 4.1 Sex differences in plaque instability and cerebrovascular symptomatology

In our study, sex was observed to be a strong determinant of plaque instability, where men exhibited more unstable plaques than women, predominantly composed of a large lipid core, intraplaque hemorrhage, and a greater presence of inflammation and cap infiltration, while women exhibited more stable plaques, which were composed predominantly of fibrous tissue. Our findings support other lines of evidence using either histopathological analyses of plaque specimens or non-invasive imaging, where clear differences in plaque morphology and composition were also demonstrated between men and women regardless of the degree of carotid artery stenosis, with men possessing higher risk carotid plaque features than women^6–8^. It is suggested that these differences in plaque composition may help partly explain why women benefit less from carotid revascularization than men (the stroke relative risk reduction following a CEA is only 4% in women compared to 51% in men) and have increased perioperative stroke risk^11, 12, 24, 25^. As a result, often in clinical practice less women are selected to undergo a CEA than men as there exists much uncertainty in deciding the appropriate management for carotid disease in women. In our own study, women comprised 30% of the total CEA population, who are consecutively recruited (acceptance rate: 99%). As evident by the baseline characteristics of our population shown in **Table 1**, this population is representative (i.e., sex distribution, comorbidities, risk factor profile) of the general population with atherosclerosis undergoing carotid intervention^26, 27^. Thus, clinical selection bias could represent a limitation, as the vascular surgeons are more likely to select, based on the current guidelines, women who they believe represent the most severe cases to be operated on and are the most likely to benefit from surgical intervention. In most cases, these are women who suffered a symptomatic event. This explains why a similar proportion of men and women were observed to be symptomatic in our study population, and experienced similar types of ischemic events, despite stroke incidence being higher among men than women in the general population. Interestingly, despite many women being symptomatic, their plaques exhibited minimal features of “plaque instability” (according to the traditional concept of the unstable plaque) compared to men. It has been suggested that thrombosis occurs via diverse mechanisms in either sex; thrombi overlying fibrotic atherosclerotic plaques in women may likely form due to surface endothelial erosion, rather than typical cap rupture, which is frequently observed in men’s unstable plaques^28^. Thrombosis in such cases are often triggered by an enhanced systemic thrombogenic state (i.e., enhanced platelet aggregability and activation) promoted by traditional risk factors, such as hypertension, smoking, and hypercholesterolemia^29^. Notably, in our study population, women with thrombotic plaques had significantly higher circulating platelet counts compared to their male counterparts, despite no differences in anti-platelet therapy.

### 4.2 Sex differences in circulating lipid, immune, and adipokine profiles

Significant sex differences observed at the level of the plaque led us to hypothesize that significant differences between men and women would also be observed at the level of the circulation. Therefore, we explored certain circulating lipid, immune, and adipokine markers that could potentially reflect and contribute to the sex-specific features in the plaque. Overall, women had a more anti-atherogenic plasma lipid profile than men. Specifically, higher concentrations of circulating HDL-C and apoA-I were detected in women compared to men. It has been shown that HDL can maintain plaque stability by inhibiting degradation of the fibrous cap extracellular matrix through its anti-elastase activity^30^, which may in part help explain why women generally have more stable plaques than men. Although these findings of higher circulating HDL-C and apoA-I among women than men have been consistently documented in other studies, particularly when comparing premenopausal women to age-matched men, the mechanism for this sex-specific difference remains not well understood^31^. It is suggested that women may have a greater HDL-apoA-I synthesis rate than men, which is associated with enhanced cholesterol efflux capacity^32^. Sex is also a biological variable that affects the functions of the immune system^33^. In our study population, men were observed to have a greater proportion of circulating monocytes compared to women, reflecting the larger abundance of macrophage accumulation observed at the level of their plaques. In previous studies, men have also been reported to have more circulating CD14 and CD16 monocytes compared to women, as well as elevated inflammatory cytokine and chemokine production^33, 34^. However, no differences in selected cytokine, chemokine, and vascular injury markers (hsCRP, IFNγ, IL-1β, IL-6, IL-10, MIP-1α, TNFα) were reported in a representative subset of our participants. Sex differences in adipose tissue biology also exist, with respect to the release of adipokines. In our study population, women had significantly higher circulating total and HMW adiponectin levels than men, as well as higher HMW to total adiponectin ratio. These findings are in line with multiple studies that have established higher circulating adiponectin levels among women compared to men^35, 36^. Adiponectin is the most abundantly secreted adipose tissue-derived protein, which exerts insulin-sensitizing, anti-inflammatory, and vasculoprotective properties. Thus, higher circulating HMW adiponectin (the biologically active form)^37, 38^ and a higher HMW to total adiponectin ratio may also partially explain why women have a lower risk of developing unstable atherosclerotic disease than men.

In addition to reporting sex differences in circulating lipid, immune, and adipokine profiles among men and women with severe carotid atherosclerotic disease, we are the first to reveal an interaction between biological sex and several of these circulating parameters on influencing plaque instability. These markers may help identify men or women with “unstable” plaque profiles who may be better suited for surgical intervention. However, future prospective studies are needed to validate the predictive value and reliability of these parameters as sex-specific markers of plaque instability. Specifically in men, low total WBC counts, and a high monocyte to WBC ratio was observed to be associated with greater plaque instability. On the other hand, in women, a decrease in the HMW to total adiponectin ratio, also referred to as the adiponectin sensitivity index, may serve as potential markers of plaque instability. In terms of basophil to WBC ratio, while a lower ratio was associated with greater plaque instability in men, the opposite was observed in women. Although basophils are scarcely investigated in the context of cardiovascular disease, Pizzolo et al. demonstrated an increased basophil blood count to be associated with an increased risk of mortality in patients with stable coronary artery disease^39^. Since basophils are being recognized as a procoagulant player in hemostasis, a regulator of platelet aggregation, and have been proposed to play a pathophysiological role in thrombosis^39^, they may act as a potential mediator of female-specific plaque instability.

### 4.3 Strengths and limitations

To the best of our knowledge, this is the first and most comprehensive study to characterize sex differences at the level of the plaque and circulation in older men and postmenopausal women with stable and unstable carotid atherosclerotic plaques, as well as examine the associations between circulating lipid, immune, and adipokine parameters of carotid plaque instability. However, our study has certain limitations. Firstly, our results may not be generalizable to the general population of men and women with carotid atherosclerosis, since our study includes a more specific population of men and women who were selected for a CEA. Therefore, as mentioned above, due to clinical selection bias, men are more often selected for carotid revascularization than women, while those women who undergo surgical intervention often represent the most severe cases, having previously suffered a cerebrovascular ischemic event. Nonetheless, women were observed to have more favorable circulating profiles and plaques with minimal features of plaque instability compared to men. Secondly, because of the cross-sectional nature of the study design, the direction of causality among the observed relationships cannot be inferred. Although this limits the establishment of the true diagnostic potential of the proposed circulating markers in predicting plaque instability, our study acts as the first phase (proof-of-concept phase) in the evaluation of novel biomarkers.

## 5. Conclusion

Our study provided much-needed evidence that sex differences exist at the level of the blood and plaque between men and women with severe carotid atherosclerotic disease. Men were observed to have less favourable circulating profiles compared to women, and more unstable plaque phenotypes. Overall, our study results place importance on the fact that the ‘one size fits all’ approach should not be applied in carotid disease management, and that there is a dire need for sex-specific guidelines. The current guidelines are lacking sex-specific orientation and are based solely on the degree of carotid artery stenosis, which has been proven to be an incomplete determinant of plaque instability. Therefore, intensified efforts to improve the screening and identification of high-risk plaques tailored to men and women are pivotal to deliver optimal individualized treatment. Herein, we identified several potential circulating markers that relate to sex-specific plaque phenotypes for better prediction of high-risk plaques in women and in men. Importantly, these markers are simple to measure and often assessed in routine examination, making them easily implementable into clinical practice. Upon future validation, the identified predictive markers could be implemented into clinical practice to monitor when the plaque becomes unstable and help better select men and women for surgical and medical management of their plaques.

## 7. Acknowledgements

We are grateful to the vascular pathologists (Drs Veinot and Lai) for their histological assessments of plaque instability. Furthermore, we are thankful to the vascular surgeons (Drs Steinmetz, MacKenzie, Corriveau, Obrand, Gill, and Bayne) at the MUHC and JGH for their help with subject recruitment.

